# Reconstructing subclonal composition and evolution from whole genome sequencing of tumors

**DOI:** 10.1101/006692

**Authors:** Amit G. Deshwar, Shankar Vembu, Christina K. Yung, Gun Ho Jang, Lincoln Stein, Quaid Morris

## Abstract

Tumors often contain multiple, genetically distinct subpopulations of cancerous cells. These so-called subclonal populations are defined by distinct somatic mutations that include point mutations such as single nucleotide variants and small indels - collectively called simple somatic mutations (SSMs) - as well as larger structural changes that result in copy number variations (CNVs). In some cases, the genotype and prevalence of these subpopulations can be reconstructed based on high-throughput, short-read sequencing of DNA in one or more tumor samples. To date, no automated SSM-based subclonal reconstructions have been attempted on WGS data; and CNV-based reconstructions are limited to tumors with two or fewer cancerous subclonal populations and with a small number of CNVs.

We describe a new automated method, PhyloWGS, that can be applied to WGS data from one or more tumor samples to perform subclonal reconstruction based on both CNVs and SSMs. PhyloWGS successfully recovers the composition of mixtures of a highly rearranged TGCA cell line when a CNV-based method fails. On WGS data with average read depth of 40 from five time-series chronic lymphocytic leukemia samples, PhyloWGS recovers the same tumor phylogeny previously reconstructed using deep targeted resequencing. To further explore the limits of WGS-based subclonal reconstruction, we ran PhyloWGS on simulated data: PhyloWGS can reliably reconstruct as many as three cancerous subpopulations based on 30-50x coverage WGS data from a single tumor sample with 10’s to 1000’s of SSMs per subpopulation. At least five cancerous subpopulations can be reconstructed if provided with read depths of 200 or more.

PhyloWGS is the first automated method that can be applied to WGS tumor data that accurately reconstructs the frequency, genotype and phylogeny of the subclonal populations based on both SSMs and CNVs. It also provides a principled, automated approach to combining overlapping SSM and CNV data. By demonstrating the utility of PhyloWGS on medium depth WGS data, including from examples with highly rearranged chromosomes, we have greatly expanded the range of tumors for which subclonal reconstruction is possible.

## 1 Introduction

Tumors often contain multiple, genetically diverse subclonal populations of cells that have evolved from a single progenitor population through successive waves of expansion and selection [1]. Analyses of this evolutionary history can help identify characteristic series of driver mutations associated with cancer development and progression [2,3]; and can provide insight into how tumors might respond to treatment [4, 5] In some cases, it is possible to genotype the subpopulations present in a tumor, as well as reconstruct their evolutionary history, using the population frequencies of mutations that distinguish these subclonal populations [6–20]. However, this reconstruction is increasingly being done through whole genome sequencing (WGS) of bulk tumor samples (see, e.g., [21]) and no automated methods yet exist that can reliably perform this reconstruction on the basis of these data.

Subclonal reconstruction algorithms attempt to infer the population structure of heterogeneous tumors based on the measured allelic frequency of their somatic mutations. Some methods perform this reconstruction based solely on point mutations (also known as ‘simple somatic mutations’ or SSMs) [15-18,20,22]. Others use changes in read coverage to identify regions with an average ploidy that differs from normal. They then attempt to explain these changes by copy number variations (CNVs) affecting some of the tumor cells [14,19,23,24].

However, because of the low read depth, WGS data presents a particular challenge to subclonal reconstruction methods. Detecting subclonal populations requires accurate estimates of the proportion of cells with each mutation (i.e., *their population frequency*). To obtain these estimates, existing SSM-based methods rely on targeted resequencing where the read depths are orders of magnitude larger than current WGS depths [16,17,22]. On the other hand, WGS reveals many more somatic mutations, and preliminary evidence suggests that the much larger number of mutations can compensate for their decreased read depth [21]. In contrast, CNVs affect large, multi-kilobase or megabase sized regions of the genome and so the average ploidy of such a region can be accurately estimated with typical WGS read depths. Unfortunately, CNV-based subclonal reconstruction is more difficult than SSM-based reconstruction because the former requires simultaneously estimating the population frequency and the new copy number. This leads to ambiguities in the reconstruction which most CNV methods resolve by only attempting to reconstruct the ‘clonal’ cancerous population [23,24] that contains the mutations shared by all of the cancerous cells. THetA [14] and SomatiCA [19] are CNV-based methods that attempt to resolve more than one cancerous subpopulation but both appear to be limited to a maximum of two cancerous subpopulations. SSM-based methods can reliably resolve many more cancerous subpopulations [15-17,22] but so far have been applied almost exclusively to data with read depths (> 1000x) that are only available via targeted resequencing. As such, the limits of automated subclonal reconstruction based on WGS samples remains to be defined.

Another open question is how to combine CNVs and SSMs in subclonal reconstruction. Subclonal reconstruction based on SSMs is an easier statistical inference problem, but CNVs are common [25] and should not be ignored. Furthermore, CNVs overlapping SSMs can interfere with SSM-based reconstruction because sequencing only provides measurements of the allele frequency of SSMs (i.e., the fraction of the loci in the sample with the variant allele); whereas reconstruction requires the *population frequency* (i.e. the proportion of cells with the mutation) and overlapping CNVs change the relationship between the allele frequency and population frequency. Although some methods attempt to model the impact of CNVs on the allele frequency of overlapping SSMs [16-18,26], these methods make unrealistic assumptions and no method incorporates the constraints that phylogeny imposes on this relationship [21].

We describe PhyloWGS, the first method designed for complete subclonal phylogenic reconstruction of both CNVs and SSMs from whole genome sequencing of bulk tumor samples. Unlike all previous methods, PhyloWGS appropriately corrects SSM population frequencies in regions overlapping CNVs and is fast enough to perform reconstruction of at least five cancerous subpopulations based on thousands of mutations. We show reconstruction problems that cannot be correctly reconstructed using previous methods. We also demonstrate that in the absence of reliable CNV estimates, it is still feasible to perform automated subclonal composition reconstruction based on SSM frequency data at typical WGS read depths (30-50x), even for highly rearranged genomes where less than 2% of the SSMs lie in regions of normal copy number.

## 2 Previous work

Complete subclonal reconstruction assigns a genotype and a population frequency to each subpopulation in the sample. This genotype can be expressed as a binary vector with one element for each mutation; each element indicates the presence (or absence) of the corresponding variant in the subpopulation. Subclonal reconstruction can thus be posed as a binary matrix factorization problem [15]; however, solving this problem without further assumptions requires at least as many tumor samples as there are subpopulations, and potentially many more, to resolve ambiguities.

Previous subclonal reconstruction methods make one (or both) of two assumptions that make it possible to resolve more subpopulations than there are samples. One assumption, the ‘infinite sites assumption’, states that each mutation only occurs once in the evolutionary history of the tumor. The other assumption, a ‘parsimony’ (also called ‘sparsity’) assumption has multiple forms but generally assumes that the number of subpopulations in the tumor is small. In the following subsections, we discuss these assumptions and their validity in greater detail.

As a first step, many subclonal reconstruction methods identify clusters of mutations with similar population frequencies; however, most do not take the further step of identifying phylogenies consistent with these frequencies. This initial step only reveals the ‘subclonal lineages’ that are present in the sample; explicit phylogenies are needed to reconstruct the genotypes of the subpopulations therein. Often methods that do not perform phylogenic reconstruction conflate these two, leading to confusion and, possibly, erroneous reconstructions. Phylogenic inference is especially important when SSMs are in regions affected by CNVs, because, as we show, knowing the phylogenic relationship of the SSM and any overlapping CNVs is necessary to accurately correct for the impact of the CNV when translating from the measured allele frequency of the SSM into the estimate of population frequency necessary to define subpopulations.

### 2.1 Subpopulations versus subclonal lineages

We use the term *subpopulation* to indicate groups of cells present in non-negligible frequency in the sample that have an identical set of the identified somatic mutations. However, a given mutation may be present in multiple subpopulations and, as such, we associate each mutation with a *subclonal lineage* that contains all of the subpopulations with the mutation. The use of ‘lineage’ implies that these subpopulations are derived from a single ancestor; this is a consequence of the ‘infinite sites assumption’ discussed in detail in the next section. Although each subclonal lineage corresponds to a subpopulation that was present at some point in the tumor development; it is not necessarily still present in the sample. Subpopulations can instead be replaced by descendant subpopulations that have acquired further mutations.

### 2.2 Assumptions made in subclonal reconstruction

A major assumption, which is well-supported theoretically and empirically, is that each SSM occurs only once in the evolutionary history of the tumor cell population. By itself, this assumption, called the ‘infinite sites assumption’, can permit complete reconstruction of multiple tumor subpopulations from a single sequenced sample. Many methods also make a ‘sparsity’ (or ‘parsimony’) assumption which states that the observed mutation frequencies should be explained with as few subpopulations as possible. Under some circumstances, this assumption is well-supported, under others it can lead to incorrect reconstructions. Without explicit representation of reconstruction uncertainty, these errors can escape detection.

**Infinite sites assumption:** Formally, the *infinite sites assumption* (ISA) [16,27,28], states that in tumors, the number of possible mutations is tiny relative to the length of the genome; and so within the same tumor, mutations never occur at the same site as existing mutations. This assumption is almost always valid for SSMs, but less clearly so for CNVs which are the consequences of (often multiple) structural rearrangements affecting large portions of the genome. As such we will assume for the time being that only SSMs are being included in the phylogenic inference.

Under this assumption, cells have an SSM if and only if they gained the SSM or are descended from the cell where the SSM appeared. A consequence of this is that the binary vectors for each subpopulation must be consistent with a ‘perfect and persistent phylogeny’ [17]: each subpopulation has all of the SSMs that its ancestors had; each SSM appears in only one subclonal lineage; and each subclonal lineage corresponds to a subtree (also known as a clade) in the phylogeny of the tumor subpopulations. Subpopulations correspond to (potentially internal) nodes in the phylogeny. Once the SSMs are assigned to their lineages, then the SSM genotype of each subpopulation can be easily reconstructed [15,16].

**Parsimony:** Another common assumption is that the number of subpopulations in the sample is much smaller than the number of mutations. One form of this assumption, that we will call *clustering, or weak parsimony*, posits that clusters of mutation population frequencies correspond to single subclonal lineages. These clusters are pervasive [11,21,22]; and likely arise due to ‘selective sweeps’ occurring during the evolution of the tumor [29,30]. These sweeps, caused by newly acquired oncogenic ‘driver’ mutation(s), increase the population frequency of both the driver mutation(s) and any other somatic ‘passenger’ mutations that have previously accumulated in the cell. The clustering assumption is widely made [16-18,21,22] and is well-supported, especially for clusters of mutations with high population frequencies where the ‘pigeonhole principle’ [21] (also known as the ‘sum rule’ [16]) implies that all SSMs in the cluster must be part of a single branch of the phylogeny.

Another assumption, which we will call the *strong parsinomy* assumption, has less support, and can lead to large errors in subclonal reconstruction. Given a set of subclonal lineages, this assumption requires these lineages be explained with the minimum number of present subpopulations [15]. For example, given three clusters of SSMs, *A*, *B*, and *C*, with estimated population frequencies of the associated lineages *ϕ_A_* = 100%, *ϕ_Β_* = 60%, and *ϕ_C_* = 40% respectively, there are two subclonal reconstructions that are equally consistent with the data. The 2-subpopulation reconstruction has a branching phylogeny in which *A* is acquired first, and *B* and *C* were then acquired independently. The subpopulation with *A* and *B* constitutes 60% of cells; and the other with *A* and *C* constitutes 40% of cells and although the subpopulation containing only *A* did exist at one point in the development of the tumor, it no longer does (i.e. it has a population frequency of 0%). The 3-subpopulation reconstruction has a chain phylogeny (i.e. a single branch) and the subpopulations consist of one containing only *A* (40% of cells), another containing *A* and *B* (20% of cells), and one containing *A*, *B*, and *C* (40% of cells).

The preference for the 2-subpopulation reconstruction is not robust to errors in the estimation of the population frequencies. Increase the population frequency of *B* by 1% and only the chain phylogeny is consistent with population frequencies because the branching phylogeny requires that the *A* population have a -1% frequency. Decrease the estimated population frequency of *B* by 1% and now the branching phylogeny assigns a population frequency of 1% to the subpopulation containing only *A* and the branching and chain phylogenies both posit three subpopulations. Addressing this issue, and robustly selecting the 2-subpopulation solution, requires setting an arbitrary parameter that balances data fit with model complexity. Failure to correctly set this parameter risks not only miscalling the number of subpopulations but making large errors in their phylogeny and genotype because, as shown above, the phylogeny (and resulting genotypes) of the subpopulations for the two reconstructions are quite different.

In contrast, clustering SSMs does not increase the risk of large errors in subclonal reconstructions, except for low frequency mutations. In the example above, if *B* cluster consisted of SSMs with population frequencies between 58% and 62%, then due to the sum rule, it would be impossible to construct a phylogeny with any pair of these mutations on separate branches of the tree (i.e. 58% + 62% = 110% > 100%). So, the only risk of binning these mutations into one lineage is that we may miss low frequency (≤ 4%) subpopulations that contain some but not all of the mutations. However, if the mutation frequencies are low, the SSMs assigned to the same cluster may actually represent two (or more) distinct groups of mutations acquired in separate branches of the phylogeny. In the example above, if the *C* cluster instead contains SSMs with population frequencies between 13% to 17%, then it is possible to build a phylogeny where some of the *C* cluster mutations occur in cells with *A* and *B* and others occur in cells only containing *A*. In other cases, it is valid to cluster low frequency mutations; say the population frequencies of *A* and *B* were 100% and 90% respectively, then a chain phylogeny is the only one consistent the *C* SSM frequencies in that range.

In summary, both parsimony assumptions can lead to inaccurate phylogenic inferences and therefore inaccurate subclonal genotype reconstruction. However, the weak parsimony assumption is safe if it can be unambiguously established that the SSMs in the cluster create a phylogenic chain. This is always true if the population frequencies of the SSMs in the cluster are greater than 50%. However, with smaller SSM frequencies, algorithms that make the weak parsimony assumption but do not construct phylogenies [17,18,22] are therefore at risk of making large reconstruction errors by clustering low frequency SSMs which belong in distinct branches of the phylogeny. Also, these algorithms may infer phantom subpopulations because there is not a one-to-one correspondence between subclonal lineages and subpopulations. Furthermore, other methods, like TRaP [15], that make strong parsimony assumptions can incorrectly select a branching phylogeny when both a branching and a chain phylogeny are consistent with the frequency data. As such, neither parsimony assumption is completely risk-free, and the strong parsimony assumption is particularly prone to making large reconstruction errors if invalid.

In cases where both branching and chain phylogenies are consistent with inferred population frequencies, we suggest that one should explicitly report what is certain and what is uncertain and not rely on a single phylogeny or reconstruction [16]. To address this issue, PhyloWGS uses a tree-structured stick-breaking process prior [31] that has a weak preference for minimizing the number of subclonal lineages so as to cluster SSM frequencies when the phylogeny permits it (i.e. we only make the weak parsimony assumption). Our Markov Chain Monte Carlo (MCMC) procedure samples phylogenies from the model posterior that are consistent with the mutation frequencies and does not rule out phylogenies that are equally consistent with the data. From this collection of samples, areas of certainty and uncertainty in the reconstruction can be derived.

### 2.3 Conditions for unambiguous reconstruction of branching

Only the chain phylogeny can be unambiguously reconstructed when given only SSM population frequencies from a single sample. Unambiguously recovering branching requires either multiple samples from the tumor [10,16]; SSMs from separate branches close enough to be spanned by a single read or read pair [21]; or making the strong parsimony assumption described above [15]. With multiple samples, one can infer that *B* and *C* are on different branches if in one sample *ϕ_β_* > *ϕ_c_* and in another sample *ϕ_β_* < *ϕ_c_*; a relationship we have called the ‘crossing rule’ [16]. With nearby SSMs, if i) one can ‘phase’ the SSMs, i.e., determine that the SSMs appear on the same chromosome based on their co-occurrence with specific alleles of nearby germline heterozygous single nucleotide polymorphisms (SNPs) and ii) these SSMs do not co-occur in reads covering both SSM loci; then one has established that they do not occur in the same cells and therefore that the SSMs are in different branches of the phylogeny.

The implication of this is that without these additional data, a chain phylogeny can always be constructed that is equally consistent with the SSM frequency data as a given branching phylogeny.

### 2.4 SSM-based reconstruction

The input to SSM-based reconstruction methods consists of read counts for variant (i.e. mutant) and reference (i.e. germline) alleles for a set of SSMs. These counts provide a measurement of the allele frequency of the SSMs which must be transformed into estimates of population frequency. CNVs overlapping SSMs complicate this transformation, but in regions of normal copy number, the infinite sites assumption implies that all SSMs on automsomes are heterozygous, so the population frequency of an SSM is simply twice its allele frequency.

Existing SSM-based methods have focused on data from deep targeted resequencing of a small number of SSMs identified in an initial pass of exome or whole genome sequencing [16,17]. This targeted resequencing generates high read-depths (10^3^−10^5^) and correspondingly precise estimates of allele frequency. Semi-automated reconstruction of tumor phylogenies [21] suggests that the larger number of SSMs found in WGS could compensate for the much lower read depths, however, to our knowledge, no automated method to do this reconstruction has been published and the limits of SSM-based reconstructions on WGS data have not been defined.

### 2.5 CNV-based approaches

In principle, subclonal reconstruction based on CNVs requires less read depth than SSM-based reconstruction because large regions of the genome are affected by these changes. This allows confident detection and quantification of changes in average ploidy at typical WGS read depths [32]. Unfortunately, CNV-based subclonal reconstruction methods face a much more challenging inference problem because population frequency and copy number must be simultaneously inferred. For example, having identified a region *c* with average ploidy *x_c_* ≠ 2, and assuming that there is only one type of CNV (i.e. all cells either have this CNV or are diploid), inferring the population frequency, *ϕ*_*c*_ of cells containing the CNV requires finding values of *ϕ*_*c*_ and *C*, the new copy number, that satisfy the following equation:

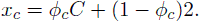

Because this equation has only one observation and two unknowns, for *x_c_* > 1 there are always at least two different solutions, even though *C* is constrained to be a non-negative integer.

In some cases, this ambiguity can be resolved using the allele frequencies of heterozygous germline SNPs, the so called ‘B-allele frequencies’, in the region affected by the CNV [21]. For example, if the observed average CN is between one and two, i.e., 1 < x_*c*_ < 2, then the equation has two solutions, either *C* = 0 and *ϕ*_*c*_ = (2 - *x_c_*)/2 or *C* = 1 and *ϕ*_*c*_ = 2 - *x_c_*. In the former case, there are an equal number of paternal and maternal copies of the CNV region in the tumor sample, so the B-allele frequencies should have a single mode at 0.5. In the latter case, there is an imbalance between the average number of copies of the paternal and maternal loci and this imbalance will manifest as a bimodal distribution in the B-allele frequencies with peaks of 1/(2 — *ϕ*_*c*_) and (1 - *ϕ*_*c*_)/(2 - *ϕ*_*c*_) (see, e.g., [21]). Although similar inferences are conceptually possible for regions where *x_c_* > 2, not all copy number changes can be resolved in this way and increased copy number generally decreases the distance between the two modes, making it difficult to distinguish unimodal and bimodal B-allele frequency distributions at high copy numbers.

If B-allele frequencies are unavailable, or cannot resolve the ambiguity, then a parsimony assumption must be employed to select *C*. Some methods only attempt to model two tumor subpopulations: the normal population and the cancerous lineage [23,24] with the highest population frequency (also known as the ‘clonal population’). These methods are useful for detecting the proportion of cells in the tumor sample that are cancerous (i.e. the cellularity), as well as detecting CNVs that are shared by all cancerous cells in the tumor, but they can fail when there are multiple subclonal populations, especially if they share few CNVs [14,19]. Recently, two CNV-based algorithms have been described that attempt to model more than one cancerous subpopulations [14,19]. These algorithms attempt to reconstruct the entire genotype of multiple subpopulations, and forego the infinite sites assumption, and so, allow different subpopulations to have different non-normal copy number. This introduces substantial ambiguity into the subclonal reconstruction; and, as such, they need to make a strong parsimony assumption in order to make the reconstruction problem statistically well-defined. As we indicated above, if invalid, the strong parsimony assumption can lead to severe reconstruction errors by inappropriately selecting branching phylogenies over chain phylogenies when both are equally consistent with the data. Furthermore, potentially due to the increased ambiguity, no CNV-based method has yet demonstrated the ability to perform reliable subclonal reconstruction from a single sample with more than two cancerous subpopulations.

### 2.6 Combining SSMs and CNVs

In loci affected by CNVs, computing the population frequency of an SSM from its allele frequency can require knowing not only the new number of maternal and paternal copies of the locus but also whether the SSM occurred before, after or independently of the CNV. This information is not always available.

On the other hand, overlapping SSMs and CNVs can provide further information about the phylogeny of the tumor. For example, if an SSM occurs in a region that is homozygously deleted in some cells (i.e., *C* = 0), then the SSM clearly cannot occur in cells containing the deletion and must either have occurred in an ancestral lineage or in a different branch of the phylogeny.

Some subclonal reconstruction methods simply ignore the impact that CNVs have on the relationship between SSM population and allele frequency [15,20]. Other methods that do account for the effect of copy number changes on SSM frequencies [16–18] integrate over all the possible relationships between allele frequency and population frequency without making use of the fact that the infinite sites assumption, which was necessary to uniquely associate SSMs to subclonal lineages in the first place, also constrains this relationship [21].

For the first time, we describe an automated method, PhyloWGS, that performs subclonal reconstruction using both CNVs and SSMs. By combining information from both CNVs and SSMs, and properly accounting for their interaction, we provide a more comprehensive and accurate description of subclonal genotype than existing methods.

## 3 Results

In the following, we first provide a brief explanation of how PhyloWGS incorporates both SSMs and CNVs in phylogenic reconstruction by converting CNVs into pseudo-SSMs and performing subclonal reconstruction on the SSMs and pseudo-SSMs; full details are provided in the Methods section. Then, we show an illustrative example where correctly accounting for the effect of CNVs on SSMs permits the correct subclonal reconstruction of a tumor population whereas using either CNV or SSM data in isolation does not. Then, we describe the results of applying PhyloWGS to simulated WGS data of different read depths, number of subpopulations, and SSMs. Next, we describe the application of PhyloWGS to three TCGA benchmark datasets. Finally, we describe the application of PhyloWGS to multiple-sample WGS data from a patient with chronic lymphocytic leukemia.

### 3.1 Incorporating CNVs with SSMs in phylogenic reconstruction

We assume that a CNV algorithm has already been applied to the sequencing data and that this algorithm provides estimates of copy number *C_i_* and population frequency *ϕ_i_* for each CNV *i*. We use these estimates in two ways: first, for each CNV, we create an equivalent pseudo-SSM with population frequency *ϕ_i_* by adding an SSM to the dataset with total reads *d_i_* and variant reads *d_i_* * *ϕ_i_*/2 rounded to the nearest whole number (this is the expected number of variant reads of a heterozygous mutation with population frequency *ϕ_i_*). We set *d_i_* to a user-defined multiple of the average WGS read-depth.

Second, we associate all SSMs within the region affected by the CNV to this pseudo-SSM. Our model (described in the Methods section) uses this association to compute the transformation between the inferred population frequency of an SSM to its expected allele frequency.

Here, we briefly describe how this transformation is done in the case that there is only one CNV affecting the SSM locus; the Methods describes the general version of the transformation that allows multiple CNVs to affect the locus. That general version is the one implemented in PhyloWGS.

Given the population frequency of the CNV, *ϕ_c_*, the copy number of the CNV *C* broken down into maternal and paternal components *C* = *C^p^ + C^m^*, and the population frequency of the SSM, *ϕ_s_*., the equations below compute the expected allele frequency of the SSM *x_ssm_*. Here we are using the terms ‘maternal’ and ‘paternal’ simply to distinguish the two copies and not to suggest that we have actually assigned each chromosome to each parent.

If a SSM lies in a region affected by a CNV, there are three possibilities for their phylogenic relationship:

1. SSM precedes the CNV event, i.e., the CNV occurs in a cell already containing the SSM
2. SSM occurs after the CNV event, i.e., the SSM occurs in a cell already containing the CNV
3. SSM and CNV occur in separate branches of phylogeny, i.e., the mutations occur in separate cells and no cell contains both the SSM and the CNV.

#### 3.1.1 Case 1 (SSM → CNV)

The description here assumes that the SSM is on the maternal strand, if it is on the paternal strand, replace *C^m^* with *C^p^* below. Also note that because of the phylogenic relationship *ϕ_s_* ≥ *ϕ_c_* > 0. So, cells with the CNV contain *C^m^* copies of the SSM and cells with the SSM but not the CNV only have one mutated copy. As such, the expected allele frequency can be written as:

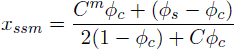

The numerator corresponds to the average number of copies per cell of the SSM-mutated locus in the population and the denominator is the average number of copies per cell of the locus (mutated or not) in the population. We note that if there is no copy number change in *C^m^* then the numerator is simply *ϕ_s_*. And if there is a copy number loss for *C^m^* then the numerator is *ϕ_s_* - *ϕ_c_*.

#### 3.1.2 Case 2 (CNV → SSM)

This case is only possible if the strand still exists after the CNV (i.e. *C^M^* ≥ 1), and the phylogenic relationship requires *ϕ_c_* ≥ *ϕ_s_* > 0. By the infinite sites assumption, only one copy of the locus is affected, so the numerator is simply *ϕ_s_* and we do not need to know the breakdown of *C* into *C^m^* and *C^p^*. As such:

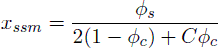

#### 3.1.3 Case 3 (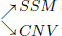)

In this case, the SSM and CNVs lie on different branches of the phylogeny and no cell in the population contains both mutations. As per Case 2, the average number of loci affected by the SSM is *ϕ_s_*. So the expected allele frequency is identical to case 2 but there is no longer an ordering of *ϕ_s_* and *ϕ_c_*:

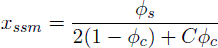

There are a few important consequences of these observations. First, we note that the breakdown of *C* into *C^m^* and *C^p^* and phasing the SSM only affects the expected variant allele frequency in Case 1. This is because it is the only case where a CNV event can affect a mutated strand. Although PhyloWGS requires the breakdown of *C* into *C^m^* and *C^p^*, we do not require the SSM to be phased and instead consider both possibilities when computing the likelihood. We do this because many SSMs cannot be phased [21].

Another consequence is that even if we are provided *x_ssm_*, *ϕ_c_, C^p^,C^m^* and the phasing of the SSM, in many cases there is still no way to distinguish between a branching or chain phylogeny. This is because the variant allele frequency is unaffected by CNV events affecting unmutated strands. For example, in Case 1, if there is no change in the copy number of the strand containing the SSM, then the transformation is identical to Case 2 and 3 and so the allele frequency cannot be used to unambiguously inferred branching. It is also easy to demonstrate that if there is a loss of the strand containing the SSM (i.e. *C^m^* = 0) then for any value of *x_ssm_* consistent with Case 3, one can assign a different *ϕ_s_* that is consistent with Case 1. However, for amplifications of the strand containing the SSM there are situations where given particular values of *x_ssm_, ϕ_c_, C^p^* and *C^m^* one can distinguish between Case 1 and Case 3. For example, given *x_ssm_* = 0.1, *ϕ_c_ = 0.4, C^m^* = 10, *C^p^* = 1, for Case 3 the inferred *ϕ_s_* is 0.56. However, if Case 1 were true the resulting inferred *ϕ_s_* would be negative and so Case 1 is not possible. This is true whenever *x_ssm_*\*(2(1-*ϕ_c_*) + *C*ϕ*_c_*) < (*C^m^* - 1)*ϕ_c_*. Outside of this particular case, for all other situations any branching phylogeny constructed using allele frequencies from a single tumor sample, one can construct a chain phylogeny that is equally consistent with the allele frequency data. This chain phylogeny adds at least one new subpopulation; so the selection of the branching phylogeny over the chain phylogeny in these cases depends solely on the strong parsimony assumption. But if a CNV can be broken down into maternal and paternal copies, the SSM can be phased, and amplification on the SSM affected strand occurs, then for some values of *x_ssm_*, branching can be unambiguously inferred.

### 3.2 The combination of CNVs with SSMs is required for accurate subclonal reconstruction

Consider a tumor where 25% of the cells are normal (no SSMs and diploid, population A), 25% come from a subpopulation with only SSMs (SSM1-4, population B) and 50% belong to a descendant subpopulation of B containing all the SSMs from B and adding new simple somatic variants (SSM5-8) and a homozygous deletion (CNV1) in the region containing SSM4, labelled population C. The evolutionary tree of this population is shown in Figure 1A. In reads sampled from this population, the expected variant allele frequencies for SSM1-3 are 37.5% (i.e. half of their population frequency) and for SSM5-8 they are 25%; however based on the rules described in the Methods section, the expected variant allele frequency of SSM4 is 25%. This is because all the copies of the genome at that position come from population A or B. Population A and B are present in equal proportions and only one copy in population B contains variant reads, so 25% of the genomes contain the variant allele. As such, methods that do not incorporate the CNV change at the SSM4 locus will likely (incorrectly) assign SSM4 to population C. Also, methods that incorporate only CNV information cannot detect the SSM-only subpopulation B.

**Figure 1:**
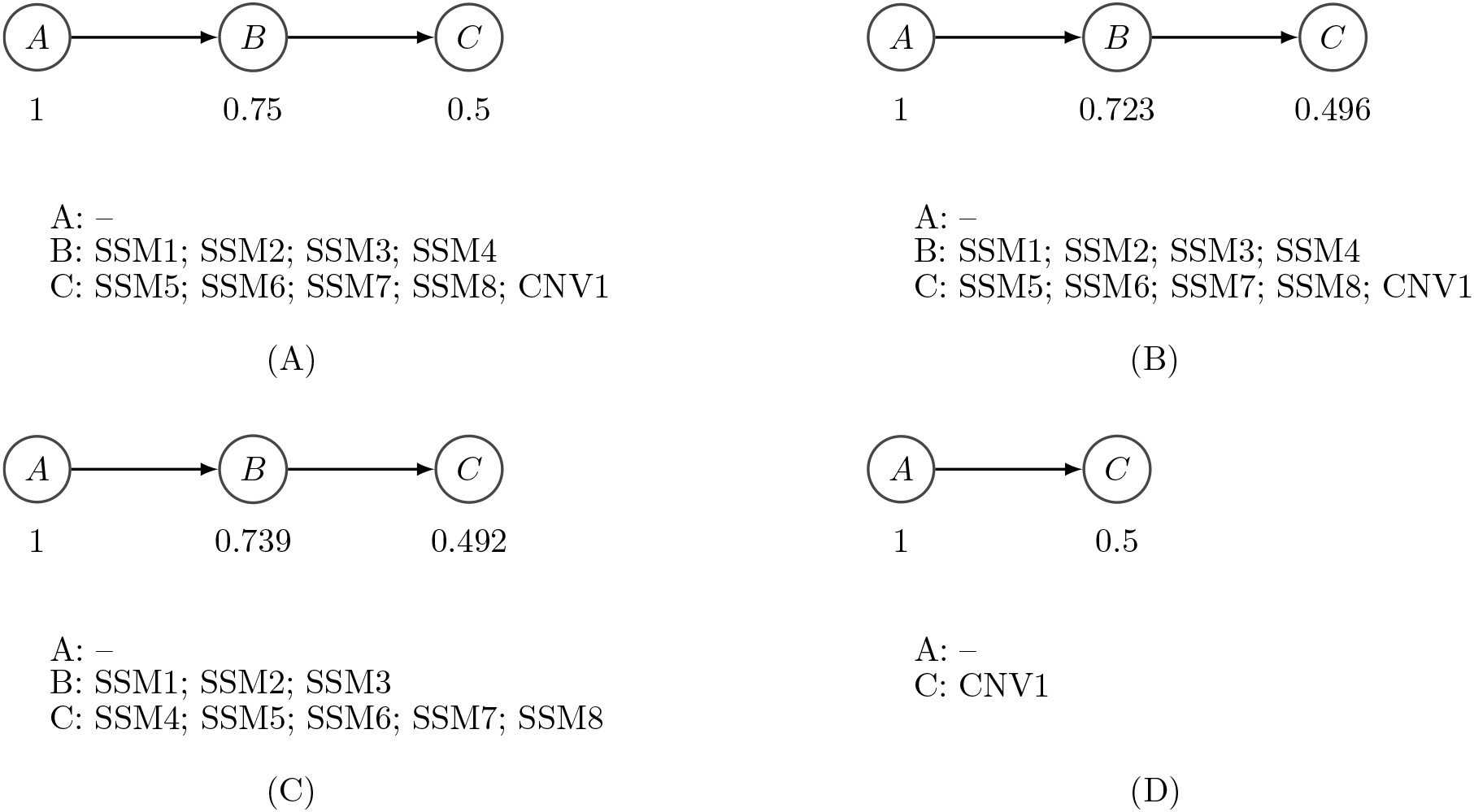
Example subclonal structure and inferred phylogenies using different methods. A: Example of tumor subclonal structure. B: Tumor phylogeny recovered by PhyloWGS. C: Tumor phylogeny recovered by PhyloSub. D: Subclonal structure implied by only CNVs.

We generated simulated variant and reference allele counts for this example at a simulated read-depth of 60. A table containing the reference and total read counts for each SSM can be found as Table 1. PhyloWGS was able to correctly reconstruct the evolutionary history and subpopulation structure (Figure 1B). However, a version of PhyloSub which ignored CNVs incorrectly assigned SSM4 to population C (Figure 1C). Furthermore, by construction, there is no way to recover population B based only on CNV data, so a perfect CNV-based algorithm would infer the subclonal structure in Figure 1D.

**Table 1:** Table showing reference and total read counts for example tumor sequencing data.

| mutation id | reference counts | total counts |
| --- | --- | --- |
| s0 | 23 | 47 |
| s1 | 35 | 69 |
| s2 | 26 | 51 |
| s3 | 29 | 56 |
| s4 | 30 | 40 |
| s5 | 42 | 70 |
| s6 | 27 | 41 |
| s7 | 36 | 56 |
| s8 | 59 | 75 |
| s9 | 57 | 76 |
| s10 | 51 | 68 |
| s11 | 37 | 51 |

Existing SSM-based methods that do not have a full model of phylogeny, such as PyClone [17], cannot accurately reconstruct the subclonal composition of this tumor because the algorithm assumes that all cancerous populations have the same number of copies at a particular locus. This logically excludes homozygous deletions. As this simple example illustrates, integrating data from both SSMs and CNVs is required for full, and accurate, subclonal reconstruction.

### 3.3 Applying PhyloWGS to simulated data

An important question in subclonal analysis of tumor samples is estimating how deep sequencing must be in order to recover the subclonal structure. To answer this question we constructed simulated read counts and applied our algorithm to this simulated data. Our simulations looked at a range of total population counts (3, 4, 5, 6), read depths (20, 30, 50, 70, 100, 200, 300) and number of SSMs per population (5, 10, 25, 50, 100, 200, 500, 1000). In each case, the first population is a normal population with no associated SSMs, while each subsequent population is a descendent of all previous populations (i.e. a chain phylogeny). For each simulated SSM *k* in population *u*, reference allele reads (*a_k_*) were drawn as:

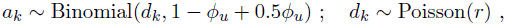

where *ϕ_u_* is the clonal frequency of population *u* and *r* is the simulated read depth. A table of the *ϕ* values used can be found as Table 2.

**Table 2:** Table of subclonal lineage proportions used.

| Number of populations | $\phi$ values used |
| --- | --- |
| 3 | 0.44, 0.11 |
| 4 | 0.56, 0.25, 0.06 |
| 5 | 0.64, 0.36, 0.16, 0.04 |
| 6 | 0.71, 0.44, 0.25, 0.11, 0.03 |

First, we examined the time needed to complete sampling as a function of the number of SSMs (shown in Figure 2). In less than one hour on a single core of an Intel Xeon E7-4830, inference could be completed as long as the total number of SSMs was less than a hundred and in less than a day if the number of SSMs was less than a thousand. The runtime depends primarily on the number of SSMs but has a smaller dependence on the inferred number of populations.

**Figure 2:**
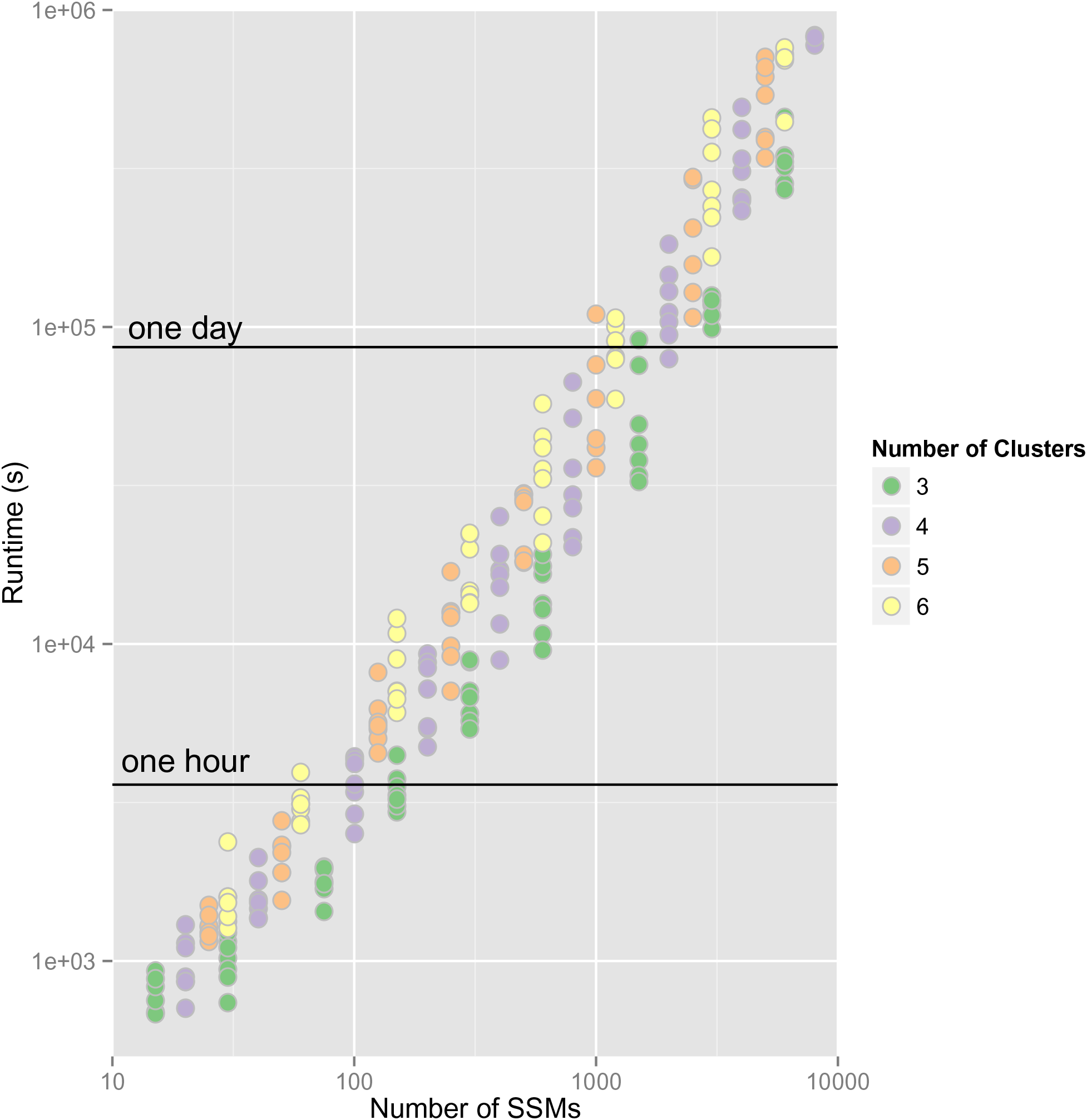
PhyloWGS Runtime. The relationship between the number of SSMs in the simulated dataset and the runtime on a log10-log10 plot. Runtime is computed using a single core of an Intel Xeon E7-4830 with 2500 MCMC iterations and 5000 inner Metropolis-Hastings iterations. Runtime can be greatly decreased by parallelizing the sampling or by taking less samples; however, the implications of these options have not been explored.

To determine the number of subpopulations our algorithm found, we analyzed the sampled tree with the highest complete data likelihood and removed any subpopulations with zero assigned SSMs. We then compared the difference between the number of subpopulations used to generate the data and the number of subpopulations identified by our algorithm. The results of this comparison are shown in Figure 3. Several relationships between simulation parameters and the output of our model can be observed. First, unsurprisingly, increasing the read depth and decreasing the number of subpopulations resulted in increased accuracy in the estimated number of subpopulations. Second, there is a U-shaped relationship between accuracy and the number of SSMs characterizing each population, where accuracy first increases and then decreases as the number of SSMs increases. This decrease in accuracy with high numbers of SSMs is non-intuitive, as more SSMs provide more information with which to perform inference. However, it is now established that the Dirichlet process prior sometimes overestimates the number of source components [33]. While this result has not been demonstrated for the tree-structured stick-breaking process used by PhyloWGS, the similarity between the processes makes it likely that this is the case. While some of these errors can be eliminated by ad-hoc removal of clusters with a small number of SSMs, there is not yet a consistent approach to do this, so we leave the results untouched. These results suggest that for three or four subpopulations, a read depth consistent with typical WGS experiments (20x-30x) is sufficient to identify the correct number of subpopulations, while experiments with 200x-300x are needed to resolve tumors with up to six subpopulations.

**Figure 3:**
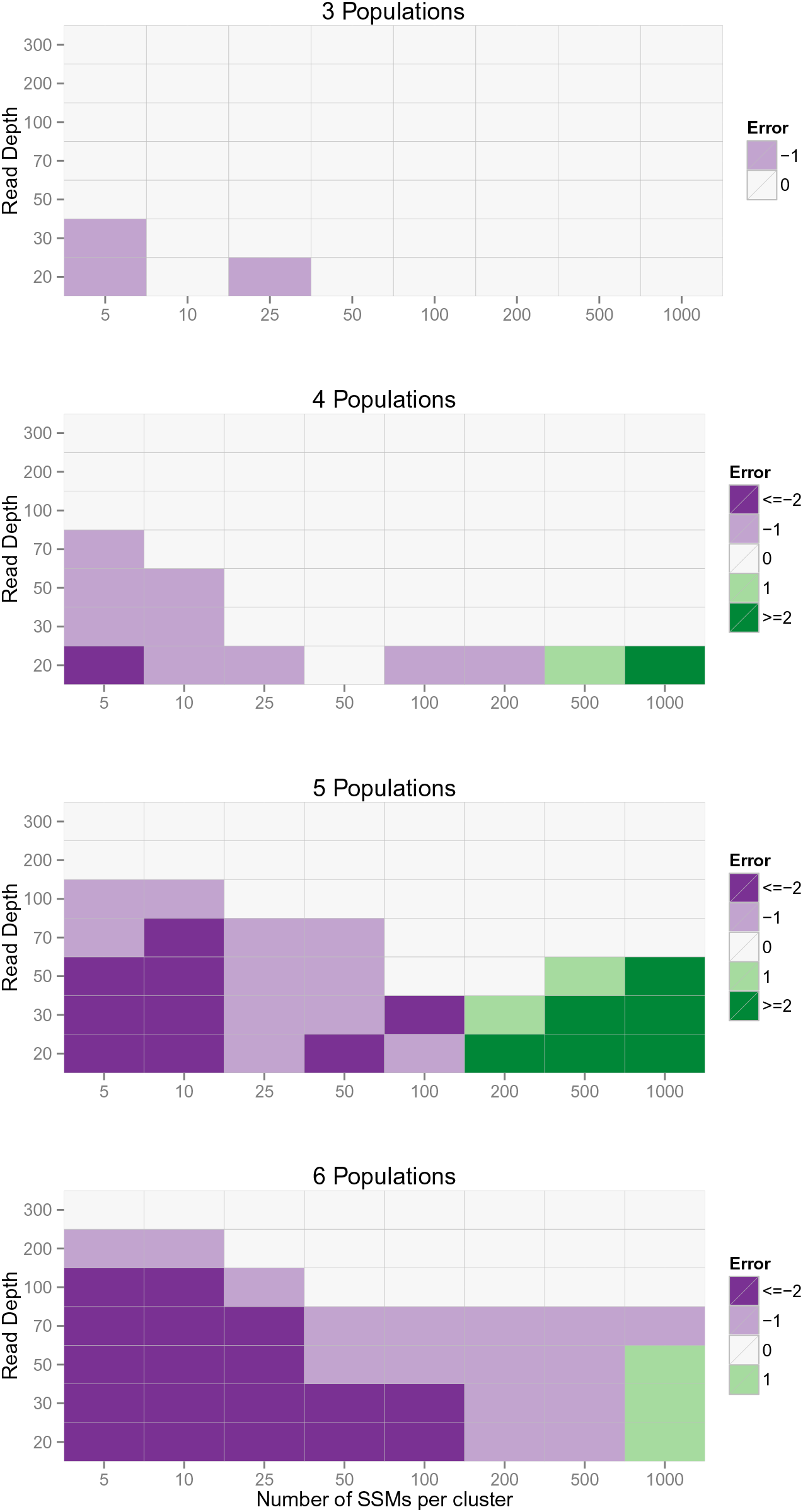
Recovering true number of clusters. Each figure shows the relationship between the number of SSMs per cluster, the read depth and the ability of PhyloWGS to recover the true number of populations for 3 (A), 4(B), 5(C) and 6(C) population simulations.

Another important measure of the performance of our algorithm is how accurately the mapping from population to SSM is. To evaluate this accuracy in a systematic way that accounts for class-imbalance, varying number of SSMs and differing number of clusters we examine the Area Under the Precision-Recall Curve (AUPRC) between the known true co-clustering matrix and the average co-clustering matrix from our samples. This average co-clustering matrix corresponds to the posterior mean co-clustering matrix and was found to better predict the true co-clustering matrix when compared to the maximum data likelihood tree. AUPRC was chosen over Area Under the Receiver-Operator Curve (AUROC) as it is known to be more informative in the presence of class-imbalance [34] which changes as the number of populations increase. The co-clustering matrix M is a binary matrix where *M_ij_* = 1 if SSM *i* and SSM *j* are in the same cluster. To provide qualitative guidance to users of the meaning of various AUPRC cutoffs, we show several examples of inferred co-clustering matrices with AUPRCs of 0.65, 0.8, 0.9 and 0.98 in Figure 4.

**Figure 4:**
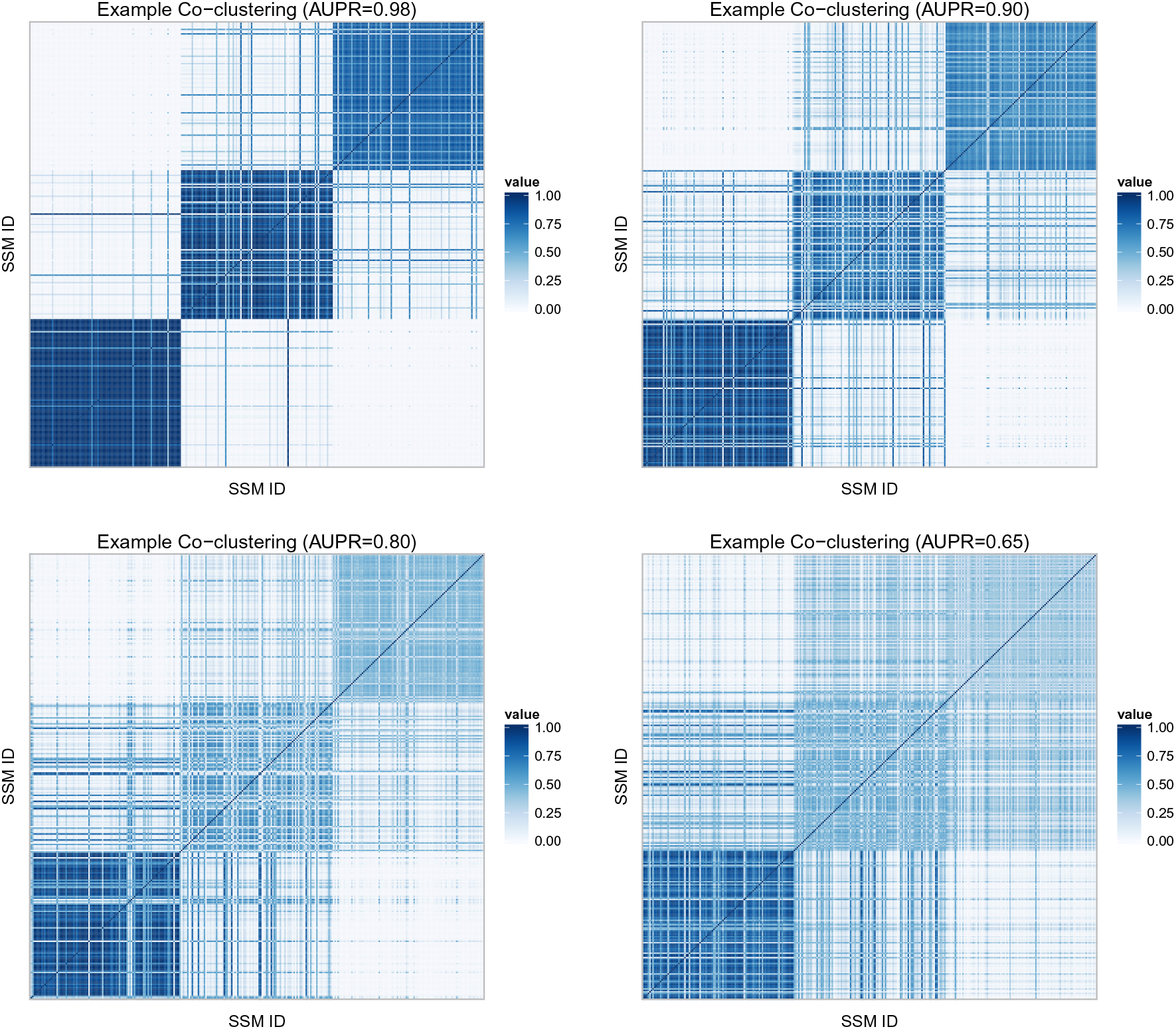
Co-clustering examples. Each figure shows mean co-clustering matrixes for simulations with 4 populations (3 cancerous), where the AUPR is 0.98 (A), 0.90 (B), 0.80 (C) and 0.6 (D). Rows and columns correspond to individual SSMs. For visibility, the matrix has been randomly subsampled to 150 SSMs from the 600 SSMs used in the simulation. Pixel color indicates coclustering probability.

In Figure 5 we plot the resulting AUPRC for our simulation experiments. As with inferring the number of populations, our method does better as the read depth increases and the number of populations decreases. Unlike the last result, there is no clear relationship between the number of SSMs and the resulting AUPRC.

**Figure 5:**
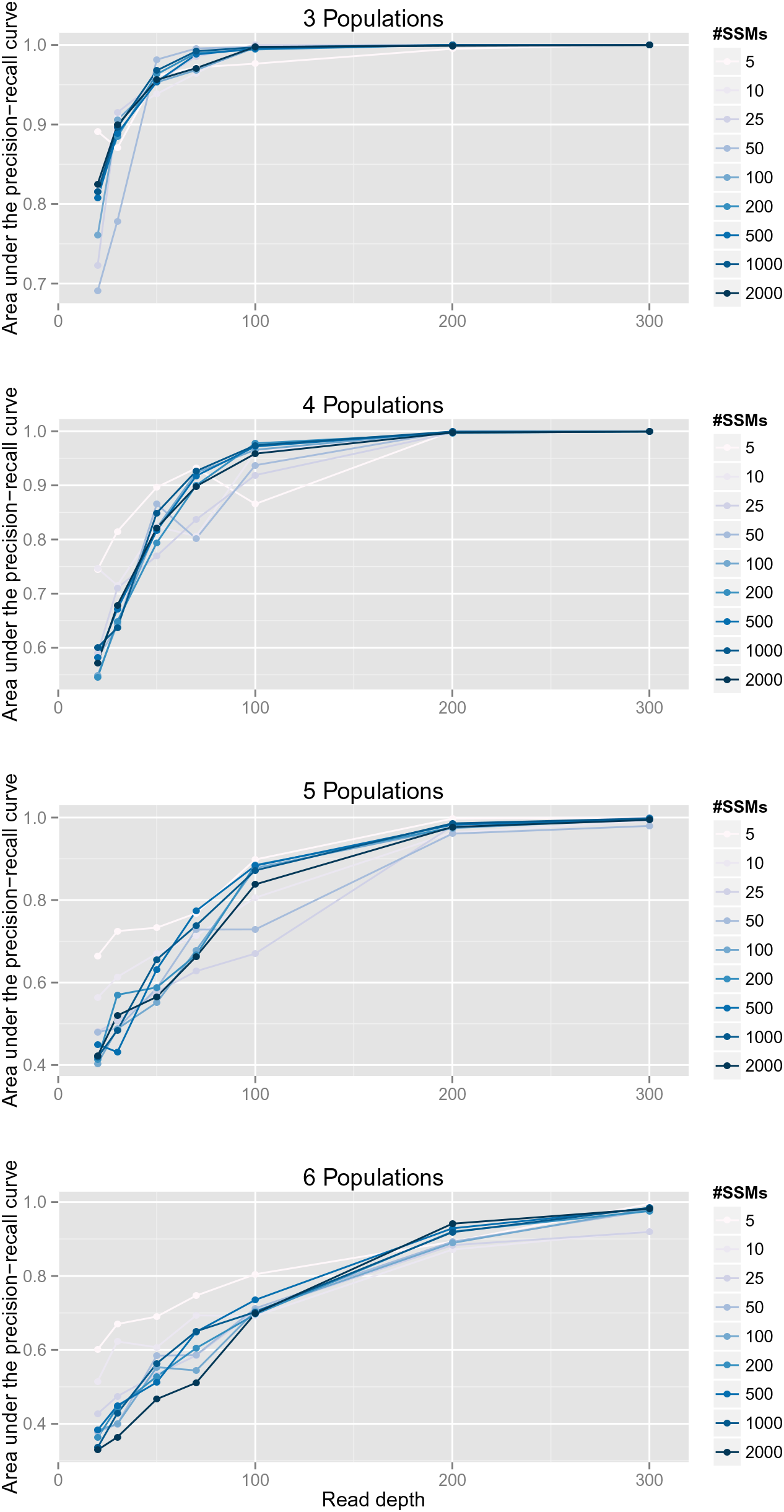
Reconstruction accuracy. Each figure shows the relationship between the read depth and the accuracy of the resulting clustering, measured as Area Under the Precision-Recall Curve (AUPR). Plots for 3 (A), 4(B), 5(C) and 6(D) populations are shown with each line representing a diifferent number of SSMs per cancerous population.

### 3.4 TCGA Benchmark

Next, we apply our algorithm to the TCGA variant-calling benchmark 4 dataset [35]. The samples we examined consist of a normal population, a cancerous cell-line population HCC 1143 and a spiked-in subclonal descendant of the cancerous population in various proportions with 30x coverage. Starting with the publicly available BAM files we identified locations of possible structural variation using BIC-seq [32] with default parameters, except for the bandwidth parameter, which was set to 1000. We changed the bandwidth parameter because we found the default value of 100 resulted in overly noisy segmentations and highly variable normalized read counts. To identify SSMs and the number of variant and reference reads for each SSM, we reverted the BAM files into unaligned reads using Picard 1.90 [36]. Reads for each sample were then realigned using BWA 0.6.2 [37] and collapsed using Picard. Aligned reads of a cancerous sample and its matched normal were analyzed by two somatic calling tools: MuTect 1.1.4 [38] and Strelka 1.0.7 [39]. A set of high confidence mutations were extracted by taking an intersection of the calls made by MuTect and Strelka. Previous verification on other tumor/normal pairs showed that this approach achieved > 90% precision (data not shown). We first ran THetA [14] using the output of BIC-seq with the aim of using THetA’s output to provide us with the CNV information that PhyloWGS requires (see Methods section). However, despite the fact that the subclonal population varied from 40% to 10%, THetA returned nearly identical composition inferences for all the samples (see Figure 6). Recognizing that we could not rely on the resulting structural variant information, we instead simply removed all SSMs in a location where BIC-seq identified possible structural variation. This eliminated most of the SSMs identified, leaving only 62 SSMs from the original 4,344. Despite this small number of SSMs our algorithm was still able to identify the correct number of populations and captured the changing composition of the samples. Also, the inferred SSM content of each cluster was identical in the three separate runs.

**Figure 6:**
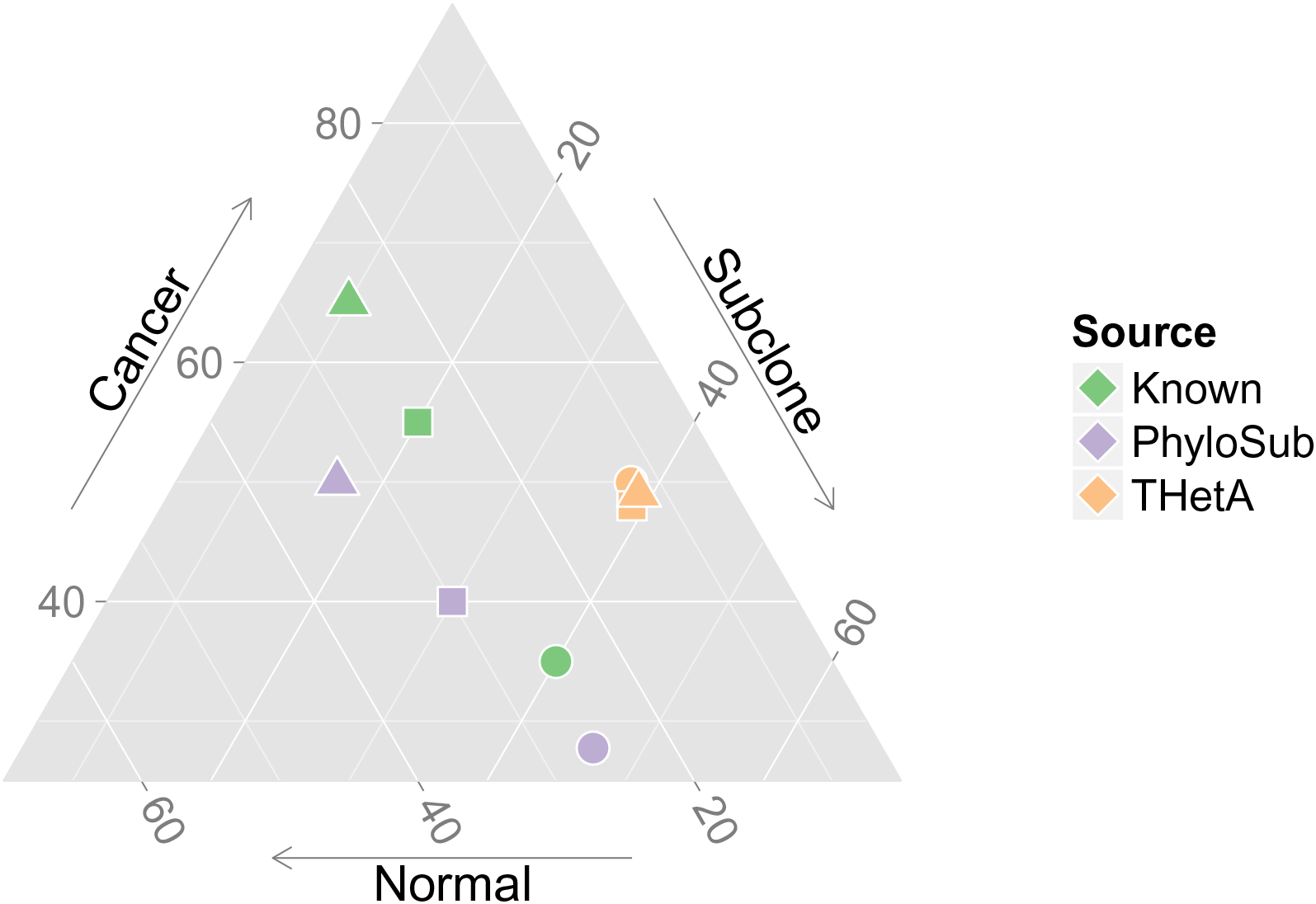
True and inferred composition of TCGA benchmark samples. The figure shows the true (red), inferred by PhyloSub (green) and inferred by THetA (orange) composition of three TCGA benchmark samples. Each point in this probability simplex corresponds to three positive proportions that sum to one.

### 3.5 Chronic Lymphocytic Leukemia

Finally, we applied PhyloWGS to a Chronic Lymphocytic Leukemia (CLL) dataset from [10]. In particular, we looked at the data from patient CLL077, found in Supplementary Table 7 of the original publication [10]. For this patient, five tumor samples were collected over the course of treatment. We note that our method does not assume or use any temporal relationships in multiple sample data and could equally be applied to multiple samples collected simultaneously. We have previously reported experiments using the targeted resequencing data with average read depth of 100,000x at 17 identified SSMs [16], we now instead use the data from WGS for that same set of mutations, with average read depth of 40x. By examining the number of reference and variant alleles it was clear that the mutation in gene SAMHD1 was at a location that was homozygous in the cancerous subpopulation it was part of. This is because the proportion of variant reads was far above 50% (the expected variant allele proportion for a heterozygous SSM present in every cell of the sample). We simulated the data that a CNA algorithm would find by assuming that the copy number at that location was 1 in a CNA-defined subpopulation and that the proportion of cells in that population was the same as implied by halving the proportion of variant alleles. After running PhyloWGS on this data, we compare the maximum data likelihood tree with the expert-generated tree found using a semi-automated method and targeted resequencing data in Figure 7. The two trees are nearly identical with the exception of assigning a single SSM to a child of the subpopulation where it is found in the expert tree.

**Figure 7:**
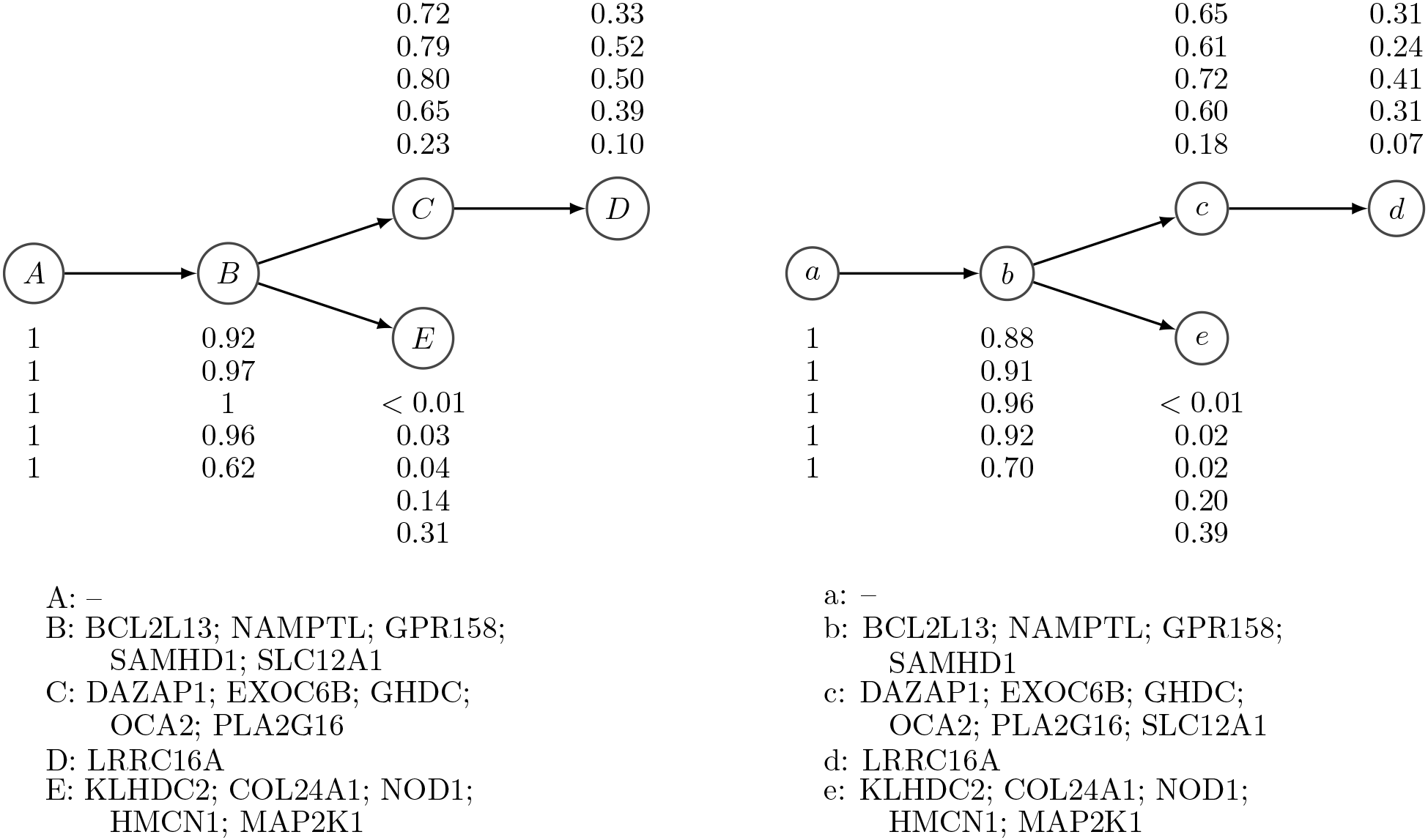
Expert-generated and inferred phylogenies from CLL patient CLL077. Left: The expert-generated phylogeny based on targeted deep sequencing data. Right: The phylogeny inferred by PhyloWGS on allele frequencies of the same SSMs found using WGS. The subclonal lineage population frequencies for the five samples and the SSM assignments of lineages are also shown in the figure.

## 4 Discussion

Our work makes two important contributions to the burgeoning field of subclonal reconstruction. First, we provide the first automated method that integrates SSM and CNV data in the reconstruction of tumor phylogenies. This is an important innovation, previous methods either ignore the impact that CNVs have on SSM allele frequencies [15,20], or assume that the CNVs affect all (and only) the cells that contain the SSM [16–18]. These assumptions can lead to incorrect inferences about the population frequency of SSMs because how a CNV affects the allele frequency of an SSM depends on its phylogenic relationship with the SSM. Many of the insights about how to integrate SSM and CNV data appear in [21]; our work here extends and formalizes these seminal observations while also providing an automated method for phylogenic reconstruction. A further advantage of combining SSMs and CNVs in the phylogenic reconstruction is that CNVs overlapping the SSM locus can provide further constraints on the tree-structure than are provided by SSM frequency alone, and we described one case where it is possible to unambiguously infer branching when an amplification of a SSM-containing locus does not lead to a large increase in the SSM allele frequency. Second, we show that given typical WGS read depths, SSM-based methods are able to accurately reconstruct tumor phylogenies and detect and assign SSMs for at least six subpopulations. Previously it wasn’t clear to what extent this reconstruction is possible; and no automated reconstructions with more than two cancerous subpopulations based on WGS data had been described. Furthermore, we report examples of accurate subclonal reconstruction for cancer populations with highly reordered chromosomes solely on the basis of SSM frequencies in the regions of normal copy number. On these same data, a state-of-the-art CNV-based method failed to perform the reconstruction.

The current version of PhyloWGS relies on preprocessing the sequencing data with a CNV-based method for subclonal reconstruction. This is because it assumes that initial population frequency *ϕ_i_* and copy number data *C_i_* are already available for the CNVs; furthermore, for amplifications, *C_i_* > 2, it requires *C_i_* to be separated into the relative number of each of the two copies, i.e., {*C_i_^m^*, *C_i_^P^*}. It does not, however, require the SSMs to be phased; in other words, it does not need to know whether the SSMs occurred on the maternal or paternal copy of the chromosome. Currently, no CNV-based automated method can provide {*C_i_^m^*, *C_i_^P^*}, so our work anticipates developments in this methodology. Until such a method is available, PhyloWGS can rely on regions of copy number loss (i.e. *C_i_* < 2), where there is only one possible breakdown. Furthermore, these estimates are only required when an SSM precedes a CNV on the same branch of a phylogeny, so SSMs in amplified regions can still be placed in the phylogeny with subpopulations that follow the CNV or are on separate branches of the phylogeny from the CNV.

We also note that PhyloWGS does not require the CNV-based preprocessing to be able to detect all of the subclonal populations, and we have shown that PhyloWGS can detect additional populations either defined completely by SSMs or that were not detected by CNV-based methods. This is particularly important because current CNV-based methods are limited to a maximum of two cancerous populations and those that can detect > 1 cancerous subpopulation do so by relying on a strong parsimony assumption. If invalid, this assumption can lead to large errors in subclonal reconstruction because it selects branching phylogenies over chain phylogenies that are equally well-supported by the data.

Indeed our results suggest an alternative strategy for combining SSMs and CNVs in subclonal reconstruction. Regions unaffected by CNVs can be relatively easily detected using methods such as BIC-seq [32]. Even in highly rearranged cancer genomes, there are often non-negligible regions of normal copy number and we have shown that we can achieve reasonably accurate subclonal reconstructions using the limited number of SSMs in regions of normal copy number in the TCGA benchmark. As such, we propose that SSM-based subclonal reconstruction should be performed first on SSMs in regions of normal copy number. This initial reconstruction can then guide the assignment of CNVs to subpopulations: the population frequencies derived from the initial SSM-based reconstruction simplify the inference of CNVs by removing one of the unknowns normally present in CNV-based reconstruction. Finally, the SSMs in CNV-affected regions can then be assigned to subpopulations. A major advantage of this approach is that it avoids the problematic strong parsimony assumption that guides current CNV-based methods. However, this approach would miss subclonal lineages defined purely by CNVs.

In the final stages of preparing this manuscript, a new method, cloneHD [26] was published. Like PhyloWGS, this method combines both SSMs and CNVs in subclonal reconstruction and does so using WGS data from single and multiple samples. However, unlike PhyloWGS, cloneHD does not explicitly perform phylogenic reconstruction, so it is unable to fully account for the phylogenic relationship among SSMs and CNVs when analyzing SSM allele frequency. As such, it is not clear to us that it can correctly solve the subclonal reconstruction problem posed in Figure 1. The cloneHD manuscript also does not extend the limits of WGS-based subclonal reconstruction as none of the examples reconstruct more than two cancerous subpopulations from a single sample. Finally, cloneHD appears to rely on the strong parsimony assumption in order to assess subclonal genotypes, and only reports a single reconstruction, obscuring the the uncertainty involved. However, cloneHD does appear to be an interesting and powerful method and we hope that future work can compare the merits and drawbacks of these alternate approaches to subclonal reconstruction.

## 5 Conclusions

We have presented a new method, PhyloWGS, that reconstructs tumor phylogenies and characterizes the subclonal populations present in a tumor sample using both SSMs and CNVs. Our method takes as input measures of allelic frequency of SSMs, as well as estimates of population frequencies and copy number for each CNV. PhyloWGS groups the SSMs and CNVs into subpopulations, and estimates the population frequencies and the phylogenic relationship of these subpopulations. PhyloWGS is based on a generative probabilistic model of allele frequencies that incorporates a nonparametric Bayesian prior over trees. The output of PhyloWGS consists of samples from this distribution generated through Markov Chain Monte Carlo and we report the tree that maximizes the likelihood of the data found during the sampling run, if a single point estimate is required. However, unlike our previous PhyloSub method [16], PhyloWGS includes CNVs in its subclonal reconstruction and, in doing so, can correctly account for the effect of CNVs on the allele frequency of overlapping SSMs. PhyloWGS also runs 50x faster than PhyloSub, making it feasible to apply it to the thousands of SSMs that are found through WGS.

We have applied PhyloWGS to real and simulated data from WGS of tumor samples to demonstrate that subclonal populations can be reliably reconstructed based solely on SSMs from medium depth sequencing (30x-50x). We have also used PhyloWGS to correctly solve a simulated subclonal reconstruction problem that neither an SSM-based nor CNV-based method can solve alone; and to reconstruct the phylogeny and subclonal composition of a highly rearranged sample for which a CNV-based method fails. Finally, using WGS of time-series samples from a chronic lymphocytic leukemia patient, PhyloWGS recovers the same tumor phylogeny previously reconstructed by applying PhyloSub and a semi-manual method to data from deep targeted resequencing.

Our work thus greatly expands the range of tumor samples for which subclonal reconstruction is possible, enabling widespread use of automated subclonal reconstruction for medium-depth WGS sequencing experiments.

## 6 Methods

### 6.1 PhyloSub model

Our probabilistic model for read count data is based on PhyloSub [16]. For each SSM that is detected by high-throughput sequencing methods, cells containing the SSM are referred to as the variant population and those without the variant as the reference population. Let *a_i_* and *b_i_* denote the number of reads matching the reference allele A and the variant allele B respectively at position *i*, and let *d_i_* = *a_i_* + *b_i_*. Let *μ_i_^r^* denote the probability of sampling a reference allele from the reference population. This value depends on the error rate of the sequencer. Let *μ_i_^v^* denote the probability of sampling a reference allele from the variant population which is set to 0.5 if there are no CNVs – in other words, the SSM is assumed to only affect one of the two chromosomal locations. Let 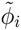 denote the fraction of cells from the variant population, i.e., the SSM population frequency at position *i*, and 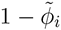 denote the fraction of cells from the reference population at position *i*. Let DP(*α*, *H*) denote the Dirichlet process (DP) prior with base distribution *H* and concentration parameter *α*. Samples from the DP are used to generate the SSM population frequencies 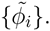 The observation model for allelic counts has the following generative process:

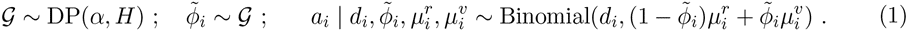

The posterior distribution of 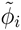 is 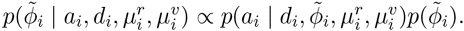

The Dirichlet process prior DP(*α*, *H*) in the observation model of allelic counts (1) is useful to infer groups of mutations that occur at the same SSM population frequency [11,17]. Furthermore, being a nonparametric prior, it allows us to avoid the problem of selecting the number of groups of mutations *a priori*. However, it cannot be used to model the clonal evolutionary structure which takes the form of a rooted tree. In order to model this, we use the tree-structured stick-breaking process prior [31] denoted by TSSB(*α*, *γ*, *H*). The parameters α and *γ* influence the height and width of the tree respectively and are similar to the concentration parameter in the Dirichlet process prior. Let 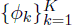 denote the set of unique frequencies in the set 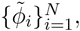 where *K* is the number of subclones or nodes in the tree. In other words, multiple elements in the set 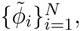 will take on the same value from the set 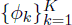 of unique frequencies. The prior/base distribution *H* of the SSM population frequencies is the uniform distribution Uniform(0,1) for the root node and Uniform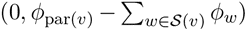 for any other node *v* in the tree, where par(*v*) denotes the parent node of *v* and *S*(*v*) is the set of siblings of *v*. This ensures that the clonal evolutionary constraints (discussed below) are satisfied when adding a new node in the tree. Given this model and a set of *N* observations 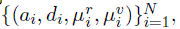 the tree structure and the SSM population frequencies 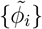 are inferred using Markov Chain Monte Carlo (MCMC) sampling (see PhyloSub [16] for further details).

Given the current state of the tree structure, we sample SSM population frequencies in such a way that the SSM population frequency *ϕ_ν_* of every non-leaf node *v* in the tree is greater than or equal to the sum of the SSM population frequencies of its children. To enforce this constraint, we introduce a set of auxiliary variables {*η_ν_*}, one for each node, that satisfy 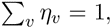 and rewrite the observation model for allelic counts (1) explicitly in terms of these variables resulting in the following posterior distribution:

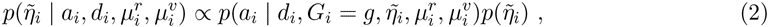

where we have used 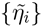 to denote the auxiliary variables for each SSM. The prior/base distribution of the auxiliary variables is defined such that it is 1 for the root node and Uniform(0, *η*_par(*ν*)_) for any other node *v* in the tree. When a new node *w* is added to the tree, we sample *η_w_* ∼ Uniform(0, *η*_par(_*_w_*_)_) and update *η*_par(_*_w_*_)_ *←η*_par(_*_w_*_)_ - *η_w_*. This ensures that 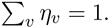 This change is crucial as it allows us to design a Markov chain that converges to the stationary distribution of {*η_ν_*}. The SSM population frequency for any node *v* can then be computed via 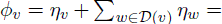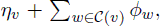 where 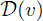 and 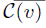 are the sets of all descendants and children of node *v* respectively. This construction ensures that the SSM population frequencies of mutations appearing at the parent node is greater than or equal to the sum of the frequencies of all its children. We use the Metropolis-Hastings algorithm [40] to sample from the posterior distribution of the auxiliary variables 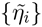 (2) and derive the SSM population frequencies from these samples by selecting the sampled set of population frequencies with the highest likelihood. We use an asymmetric Dirichlet distribution as the proposal distribution.

### 6.2 Integrating CNV data into PhyloSub

The focus of our new method, PhyloWGS is integrating SSM frequencies with existing copy number variation (CNV) based subclonal reconstructions. As mentioned above, our algorithm takes as input a set of SSMs along with their allele frequencies, expressed for each SSM *i*, as the number of reads at the locus supporting either the SSM (*b_i_*) or the reference allele (*a_i_*). We also allow our algorithm to take a set of inferred copy number changes, where for each change *j*, the input provides the new copy number *C_j_* as well as the proportion of the population with the change 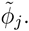 In some cases, we also require the breakdown of *C_j_* into the new number of maternal (*C_j_^m^*) and paternal (*C_j_^p^*) copies of the locus (see below for details). If this breakdown is not available, we can restrict our attention to CNVs for which *C_j_* < 2 because in these cases, there is only one possible breakdown. Also, in the absence of paternal/maternal breakdown, we should still be able to, in theory, assign SSMs with overlapping CNVs with *C_j_* > 2 to specific populations once the phylogeny and subclonal populations have been defined using SSMs and CNVs in regions of *C_j_* ≤ 2.

Below, we describe the rules, based on the infinite sites assumption, that we use to determine the relationship between the population frequency of an SSM 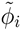 and its observed variant allele frequency (*b_i_*/*d_i_*). When the SSM does not overlap a region that has a predicted CNV in any cell in the tumor population, then the predicted allele frequency is simply half of the modeled population frequency. We also describe the method by which we transform each CNV *j* into a pseudo-SSM to be included in the phylogeny.

### If CNVs do not overlap with any SSM

If a CNV occurs in a region without any SSMs, we generate a ‘pseudo-SSM’ for the CNV *j* which is represented in the model as a heterozygous, binary somatic mutation with a read depth that reflects the uncertainty in the provided population frequency 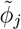 for the CNV. Specifically, we generate reference and variant read counts, *a_j_* and *b_j_*, respectively, such that the allelic frequency *b_j_*/(*a_j_* + *b_j_*) is approximately equal to *ϕ_j_*/2 and the total number of reads *a_j_* + *b_j_* is selected based on the evidence supporting the CNV. Generating this pseudo-SSM allows the CNV to be treated like any other SSM in the phylogeny model.

#### If CNVs overlap with SSMs

If a structural variant occurs in a region with an SSM *i*, this complicates the relationship between the proportion of cells that contain the SSM and the expected number of reads because cells with the CNV will have more (or fewer) than two copies of the locus where the SSM lies. Assuming equal sampling of these regions, the expected proportion of reads without the mutation ( ζ*_i_*) is always:

*N_i_^r^*/(*N_i_^r^* + *N_i_^v^*) where *N_i_^r^* is the number of copies of the locus that have the reference allele and *N_i_^v^* is the number of copies of the locus with the variant allele. To account for sequencing error we define *ϵ* as the probability of reading the reference allele when the locus contains the variant allele and vice-versa. The expected proportion of reads containing the reference allele is then:

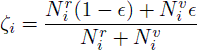

Looking at a tumor sample with multiple populations and without structural variations, if each population *u* is present with proportion *η_u_*, and where *s_i_^u^* is 1 if population *u* contains the SSM *i* and 0 otherwise, then 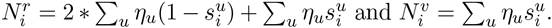 This is equivalent to an algorithm that looks at each population and performs the following update. If the population u contains the SSM *i* then

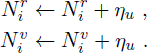

If the population does not contain the SSM then:

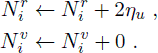

To take into account CNVs requires a more complex procedure. For each population, for each SSM, the number of reference and variant alleles depends on the copy number of the locus *C_i_* and, potentially, number of maternal (*C*_*i*_*^m^*) and paternal (*C*_*i*_*^p^*) copies of the locus *as well as* the evolutionary relationship between the SSM and the CNV. The infinite sites assumption does not apply for CNVs, adding a further level of complexity because multiple CNVs at the same locus are possible. For each population, the CNV that affects its contribution to the number of reference and variant genomes can be found by ascending the evolutionary tree towards the root. The first CNV found in this ascent is the CNV relevant for the population. If no CNV is found than the population is not affected by a CNV. For each population there are five possible situations:

i. The population does not contain the SSM and is not affected by a CNV
ii. The population does not contain the SSM but is affected by a CNV
iii. The population contains the SSM but is not affected by a CNV
iv. The population contains the SSM and is affected by a CNV, and the SSM occurred after the CNV
v. The population contains the SSM and is affected by a CNV, and the CNV occurred after the SSM

If a population does not contain the SSM, then even if a CN change has occurred (cases i and ii), the update rule is:

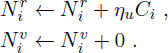

If a population contains the SSM and the SSM occurred after a CN change (or there was no CN change) (cases iii and iv) then there is a single copy of the mutated genome and the remainder are reference, so the the update rule is:

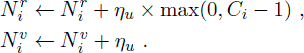

If a population contains the SSM and the SSM occurred before the CN change (case v) then there are two possibilities, the SSM is on the maternal copy or the paternal copy. If the SSM is on the maternal copy, the update rule is:

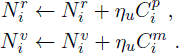

If however, the SSM is on the maternal copy, the update rule is:

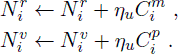

Note that the breakdown of *C_i_* into *C*_*i*_*^m^* and *C*_*i*_*^p^* is only required if the CNV occurs after the SSM on the same branch.

Now that we can calculate *N_i_^r^* and *N_i_^v^,* the observation model for the allelic counts has the following generative process (cf. (1)):

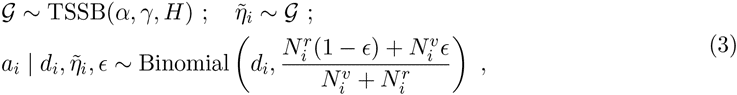

Note that in some circumstances, a SSM can be placed on a particular copy of the chromosome by looking for reads that cover the SSM and nearby heterozygous germline mutations. If this is not possible then the likelihood of *a_i_* is the average of two likelihoods; the likelihood if the SSM occurs on the maternal genome and the likelihood if the SSM occurs on the paternal genome.

### 6.3 Extension to multiple samples

Our model can be easily extended to multiple tumor samples. We make no assumptions regarding the time that the samples were collected, so this extension is equally applicable to multiple samples collected simultaneously (e.g. as in [9]) or over a period of time as in [10]. We allow the tree-structured stick-breaking process prior to be shared across all the samples. The main technical difference between the single and the multiple sample models lies in the sampling procedure for SSM population frequencies. In the multiple sample model, we ensure that the clonal evolutionary constraints are satisfied separately for each tumor sample and then make a global Metropolis-Hastings move based on the product of posterior distributions across all the samples (cf. (2)).

### 6.4 MCMC settings

In all the MCMC experiments, we fix the number of MCMC iterations to 2,500 and use a burn-in of 100 samples. We also fix the number of iterations in the Metropolis-Hastings algorithm to 5,000 and set the scaling factor for the Dirichlet proposal distribution to 100 (see PhyloSub paper [16]). We use the CODA R package [41] for MCMC diagnostics to monitor the convergence of the samplers using the complete-data log likelihood traces and the corresponding autocorrelation function.

## References

1. Peter C Nowell. The clonal evolution of tumor cell populations. Science, 194(4260):23–28, 1976.

2. D. Hanahan and R. A. Weinberg. The hallmarks of cancer. Cell, 100(1):57–70, 2000.

3. D. Hanahan and R. A. Weinberg. Hallmarks of cancer: The next generation. Cell, 144(5):646–674, 2011.

4. Samuel Aparicio and Carlos Caldas. The implications of clonal genome evolution for cancer medicine. New England Journal of Medicine, 368(9):842–851, 2013.

5. Philippe L Bedard, Aaron R Hansen, Mark J Ratain, and Lillian L Siu. Tumour heterogeneity in the clinic. Nature, 501(7467):355–364, 2013.

6. Charles G. Mullighan, Letha A. Phillips, Xiaoping Su, Jing Ma, Christopher B. Miller, Sheila A. Shurtleff, and James R. Downing. Genomic analysis of the clonal origins of relapsed acute lymphoblastic leukemia. Science, 322(5906):1377–1380, 2008.

7. N. E. Navin and J. Hicks. Tracing the tumor lineage. Molecular Oncology, 4(3):267–283, 2010.

8. A. Marusyk and K. Polyak. Tumor heterogeneity: Causes and consequences. Biochimica et Biophysica Acta, 1805(1):105–117, 2010.

9. Marco Gerlinger, Andrew J. Rowan, Stuart Horswell, James Larkin, David Endesfelder, Eva Gronroos, Pierre Martinez, Nicholas Matthews, Aengus Stewart, Patrick Tarpey, Ignacio Varela, Benjamin Phillimore, Sharmin Begum, Neil Q. McDonald, Adam Butler, David Jones, Keiran Raine, Calli Latimer, Claudio R. Santos, Mahrokh Nohadani, Aron C. Eklund, Bradley Spencer-Dene, Graham Clark, Lisa Pickering, Gordon Stamp, Martin Gore, Zoltan Szallasi, Julian Downward, P. Andrew Futreal, and Charles Swanton. Intratumor heterogeneity and branched evolution revealed by multiregion sequencing. The New England Journal of Medicine, 366(10):883–892, 2012.

10. Anna Schuh, Jennifer Becq, Sean Humphray, Adrian Alexa, Adam Burns, Ruth Clifford, Stephan M. Feller, Russell Grocock, Shirley Henderson, Irina Khrebtukova, Zoya Kingsbury, Shujun Luo, David McBride, Lisa Murray, Toshi Menju, Adele Timbs, Mark Ross, Jenny Taylor, and David Bentley. Monitoring chronic lymphocytic leukemia progression by whole genome sequencing reveals heterogeneous clonal evolution patterns. Blood, 120(20):4191–4196, 2012.

11. Sohrab P. Shah, Andrew Roth, Rodrigo Goya, Arusha Oloumi, Gavin Ha, Yongjun Zhao, Gulisa Turashvili, Jiarui Ding, Kane Tse, Gholamreza Haffari, Ali Bashashati, Leah M. Prentice, Jaswinder Khattra, Angela Burleigh, Damian Yap, Virginie Bernard, Andrew McPherson, Karey Shumansky, Anamaria Crisan, Ryan Giuliany, Alireza Heravi-Moussavi, Jamie Rosner, Daniel Lai, Inanc Birol, Richard Varhol, Angela Tam, Noreen Dhalla, Thomas Zeng, Kevin Ma, Simon K. Chan, Malachi Griffith, Annie Moradian, S.-W. Grace Cheng, Gregg B. Morin, Peter Watson, Karen Gelmon, Stephen Chia, Suet-Feung Chin, Christina Curtis, Oscar M. Rueda, Paul D. Pharoah, Sambasivarao Damaraju, John Mackey, Kelly Hoon, Timothy Harkins, Vasisht Tadigotla, Mahvash Sigaroudinia, Philippe Gascard, Thea Tlsty, Joseph F. Costello, Irmtraud M. Meyer, Connie J. Eaves, Wyeth W. Wasserman, Steven Jones, David Huntsman, Martin Hirst, Carlos Caldas, Marco A. Marra, and Samuel Aparicio. The clonal and mutational evolution spectrum of primary triple-negative breast cancers. Nature, 486(7403):617–656, 2012.

12. Scott L Carter, Kristian Cibulskis, Elena Helman, Aaron McKenna, Hui Shen, Travis Zack, Peter W. Laird, Robert C. Onofrio, Wendy Winckler, Barbara A. Weir, Rameen Beroukhim, David Pellman, Douglas A. Levine, Eric S. Lander, Matthew Meyerson, and Gad Getz. Absolute quantification of somatic DNA alterations in human cancer. Nature Biotechnology, 30(5):413–421, 2012.

13. Dan A. Landau, Scott L. Carter, Petar Stojanov, Aaron McKenna, Kristen Stevenson, Michael S. Lawrence, Carrie Sougnez, Chip Stewart, Andrey Sivachenko, Lili Wang, Youzhong Wan, Wandi Zhang, Sachet A. Shukla, Alexander Vartanov, Stacey M. Fernandes, Gordon Saksena, Kristian Cibulskis, Bethany Tesar, Stacey Gabriel, Nir Hacohen, Matthew Meyerson, Eric S. Lander, Donna Neuberg, Jennifer R. Brown, Gad Getz, and Catherine J. Wu. Evolution and impact of subclonal mutations in chronic lymphocytic leukemia. Cell, 152(4):714–726, 2013.

14. L Oesper, A Mahmoody, and BJ Raphael. Theta: Inferring intra-tumor heterogeneity from high-throughput dna sequencing data. Genome Biology, 14:R80, 2013.

15. F Strino, F Parisi, M Micsinai, and Y Kluger. Trap: A tree approach for fingerprinting subclonal tumor composition. Nucleic Acids Research, 41(17):e165, 2013.

16. W Jiao, S Vembu, A G Deshwar, L Stein, and Q Morris. Inferring clonal evolution of tumors from single nucleotide somatic mutations. BMC Bioinformatics, 15:35, 2014.

17. Andrew Roth, Jaswinder Khattra, Damian Yap, Adrian Wan, Emma Laks, Justina Biele, Gavin Ha, Samuel Aparicio, Alexandre Bouchard-Côté, and Sohrab P Shah. Pyclone: Statistical inference of clonal population structure in cancer. Nature Methods, 11:396–398, 2014.

18. Noemi Andor, Julie V Harness, Sabine Müller, Hans W Mewes, and Claudia Petritsch. Expands: expanding ploidy and allele frequency on nested subpopulations. Bioinformatics, 30(1):50–60, 2014.

19. Mengjie Chen, Murat Gunel, and Hongyu Zhao. Somatica: identifying, characterizing and quantifying somatic copy number aberrations from cancer genome sequencing data. PloS one, 8(11):e78143, 2013.

20. Nicholas B Larson and Brooke L Fridley. Purbayes: estimating tumor cellularity and subclonality in next-generation sequencing data. Bioinformatics, 29(15):1888–1889, 2013.

21. Serena Nik-Zainal, Peter Van Loo, David C. Wedge, Ludmil B. Alexandrov, Christopher D. Greenman, King Wai Lau, Keiran Raine, David Jones, John Marshall, Manasa Ramakrishna, Adam Shlien, Susanna L. Cooke, Jonathan Hinton, Andrew Menzies, Lucy A. Stebbings, Catherine Leroy, Mingming Jia, Richard Rance, Laura J. Mudie, Stephen J. Gamble, Philip J. Stephens, Stuart McLaren, Patrick S. Tarpey, Elli Papaemmanuil, Helen R. Davies, Ignacio Varela, David J. McBride, Graham R. Bignell, Kenric Leung, Adam P. Butler, Jon W. Teague, Sancha Martin, Goran Jönsson, Odette Mariani, Sandrine Boyault, Penelope Miron, Aquila Fatima, Anita Langerød, Samuel A.J.R. Aparicio, Andrew Tutt, Anieta M. Sieuwerts, Åke Borg, Gilles Thomas, Anne Vincent Salomon, Andrea L. Richardson, Anne-Lise Børresen-Dale, P. Andrew Futreal, Michael R. Stratton, and Peter J. Campbell. The life history of 21 breast cancers. Cell, 149:994–1007, 2012.

22. Li Ding, Timothy J Ley, David E Larson, Christopher A Miller, Daniel C Koboldt, John S Welch, Julie K Ritchey, Margaret A Young, Tamara Lamprecht, Michael D McLellan, et al. Clonal evolution in relapsed acute myeloid leukaemia revealed by whole-genome sequencing. Nature, 481(7382):506–510, 2012.

23. Scott L Carter, Kristian Cibulskis, Elena Helman, Aaron McKenna, Hui Shen, Travis Zack, Peter W Laird, Robert C Onofrio, Wendy Winckler, Barbara A Weir, et al. Absolute quantification of somatic dna alterations in human cancer. Nature biotechnology, 30(5):413–421, 2012.

24. Arief Gusnanto, Henry M Wood, Yudi Pawitan, Pamela Rabbitts, and Stefano Berri. Correcting for cancer genome size and tumour cell content enables better estimation of copy number alterations from next-generation sequence data. Bioinformatics, 28(1):40–47, 2012.

25. Giovanni Ciriello, Martin L Miller, Bulent Arman Aksoy, Yasin Senbabaoglu, Nikolaus Schultz, and Chris Sander. Emerging landscape of oncogenic signatures across human cancers. Nature genetics, 45(10):1127–1133, 2013.

26. Andrej Fischer, Ignacio Vázquez-García, Christopher JR Illingworth, and Ville Mustonen. High-definition reconstruction of clonal composition in cancer. Cell Reports, 2014.

27. Motoo Kimura. The number of heterozygous nucleotide sites maintained in a finite population due to steady flux of mutations. Genetics, 61(4):893, 1969.

28. Richard R Hudson. Properties of a neutral allele model with intragenic recombination. Theoretical population biology, 23(2):183–201, 1983.

29. Rebecca A Burrell, Nicholas McGranahan, Jiri Bartek, and Charles Swanton. The causes and consequences of genetic heterogeneity in cancer evolution. Nature, 501(7467):338–345, 2013.

30. Christoph A Klein. Selection and adaptation during metastatic cancer progression. Nature, 501(7467):365–372, 2013.

31. Ryan Prescott Adams, Zoubin Ghahramani, and Michael I. Jordan. Tree-structured stick breaking for hierarchical data. In Advances in Neural Information Processing Systems 23, pages 19–27, 2010.

32. Ruibin Xi, Angela G Hadjipanayis, Lovelace J Luquette, Tae-Min Kim, Eunjung Lee, Jianhua Zhang, Mark D Johnson, Donna M Muzny, David A Wheeler, Richard A Gibbs, et al. Copy number variation detection in whole-genome sequencing data using the bayesian information criterion. Proceedings of the National Academy of Sciences, 108(46):E1128–E1136, 2011.

33. Jeffrey W Miller and Matthew T Harrison. A simple example of Dirichlet process mixture inconsistency for the number of components. In Advances in Neural Information Processing Systems, pages 199–206, 2013.

34. Jesse Davis and Mark Goadrich. The relationship between precision-recall and roc curves. In Proceedings of the 23rd international conference on Machine learning, pages 233–240. ACM, 2006.

35. Adam Ewing. Tcga mutation/variation calling benchmark 4. https://cghub.ucsc.edu/datasets/benchmark\_download.html, January 2013.

36. The Broad Institute. Picard: Java tools for manipulating bam files. http://picard.sourceforge.net/.

37. Heng Li and Richard Durbin. Fast and accurate long-read alignment with burrows-wheeler transform. Bioinformatics, 26(5):589–595, 2010.

38. Kristian Cibulskis, Michael S Lawrence, Scott L Carter, Andrey Sivachenko, David Jaffe, Carrie Sougnez, Stacey Gabriel, Matthew Meyerson, Eric S Lander, and Gad Getz. Sensitive detection of somatic point mutations in impure and heterogeneous cancer samples. Nature biotechnology, 31(3):213–219, 2013.

39. Christopher T Saunders, Wendy SW Wong, Sajani Swamy, Jennifer Becq, Lisa J Murray, and R Keira Cheetham. Strelka: Accurate somatic small-variant calling from sequenced tumor-normal sample pairs. Bioinformatics, 28(14):1811–1817, 2012.

40. W Keith Hastings. Monte carlo sampling methods using markov chains and their applications. Biometrika, 57(1):97–109, 1970.

41. Martyn Plummer, Nicky Best, Kate Cowles, and Karen Vines. CODA: Convergence diagnosis and output analysis for MCMC. R News, 6(1):7–11, 2006.

